# Multimodal Evidence for Hippocampal Engagement and Modulation by Functional Connectivity-Guided Parietal TMS

**DOI:** 10.1101/2025.06.08.658503

**Authors:** Zhuoran Li, Nicholas T. Trapp, Joel Bruss, Xianqing Liu, Kang Wu, Ziyan Chen, Amit Etkin, Matthew A. Howard, Aaron D. Boes, Jing Jiang

## Abstract

Hippocampal activity supports memory and many other brain functions. Transcranial magnetic stimulation (TMS) guided by hippocampal functional connectivity (FC) shows promise in improving memory, but direct neural evidence of its capacity to engage and modulate hippocampal activity is lacking. Here we combined TMS with intracranial electroencephalography (iEEG) in 8 neurosurgical patients and with functional magnetic resonance imaging (fMRI) in 79 neurologically healthy participants. We identified that (1) single-pulse TMS to individualized parietal cortex guided by hippocampal-FC preferentially evoked distinct temporal and spectral activity patterns in the hippocampus, (2) variability in TMS-evoked hippocampal responses related to individual differences in parietal-hippocampus FC strength, and (3) repetitive TMS to hippocampal-FC-guided parietal cortex selectively suppressed hippocampal theta oscillations. These findings provide multimodal causal neural evidence and important mechanistic insights supporting the development of personalized neuromodulation strategies aimed at improving hippocampus-dependent functions.

## Introduction

The hippocampus is a deep brain structure essential for memory, emotion processing, and several other functions. Modulating hippocampal activity holds significant potential for advancing our understanding of the causal neural mechanisms underlying these functions and for developing clinical interventions for neurological and neuropsychiatric conditions linked to hippocampal dysfunction, such as Alzheimer’s disease and post-traumatic stress disorder ^1, 2^. Traditionally, hippocampal modulation has relied on invasive methods, such as deep brain stimulation ^3, 4^, limiting its broader applicability in both research and clinical settings. Noninvasive brain stimulations, such as transcranial magnetic stimulation (TMS), offer a promising alternative. Although these methods have conventionally been used to target superficial cortical regions, accumulating evidence indicates that the neural effects of TMS can propagate to remote brain regions that are structurally or functionally connected to the stimulation region, thereby influencing related cognitive functions ^5-7^. However, it remains unclear whether cortical TMS can causally engage and modulate deep hippocampal activity.

The parietal cortex, a key node within the hippocampal-cortical network, exhibits both functional and structural connectivity to the hippocampus ^8, 9^, making it a promising target for cortical stimulation to modulate the hippocampus and associated functions. Prior studies have shown that repetitive TMS (rTMS) to the parietal cortex, guided by individualized functional connectivity (FC) to the hippocampus using resting-state functional MRI (fMRI), can improve performance on hippocampus-dependent tasks, such as episodic memory ^5, 6^. These behavioral improvements were accompanied by increased parietal-hippocampal connectivity ^5, 6^. However, such connectivity changes may primarily reflect alterations of local activity at the stimulated cortical region ^10^, rather than that in the targeted hippocampus. Therefore, it remains an open question whether and how individualized FC-guided parietal TMS can effectively influence hippocampal activity. Behavioral improvements from TMS in individuals have been linked to both intrinsic FC patterns between the stimulation site and target region ^7^, and TMS-evoked response changes in deep target regions guided by targeted FC ^11^. It is therefore plausible that the extent of hippocampal engagement by parietal TMS may depend on an individual’s intrinsic FC pattern between the TMS site and hippocampus. Yet, this hypothesis has not been directly tested. Answer to this question would represent a critical prerequisite for personalized neuromodulation aiming to improve hippocampus-dependent functions.

A central challenge in addressing this knowledge gap lies in the difficulty of measuring immediate neural effects of TMS in distant subcortical regions following cortical stimulation. Recent advances in concurrent brain stimulation-recording methods, such as TMS combined with intracranial EEG (iEEG) ^12-14^ or fMRI ^15^, offer powerful solutions. For instance, our team recently developed a single-pulse TMS (spTMS)-iEEG approach in neurosurgical patients, demonstrating that spTMS targeting the dorsolateral prefrontal cortex (DLPFC) elicits rapid neural responses in temporal or spectral domains in deep brain regions, including the insula, subgenual cingulate cortex (sgACC) and dorsal anterior cingulate cortex ^12-14^. This approach is well-positioned to assess hippocampal responses to FC-guided parietal TMS and characterize its temporal and spectral dynamics. To construct a more comprehensive picture of hippocampal engagement by parietal TMS and overcome clinical constraints of iEEG exclusively allowed in neurosurgical patients, the spTMS-iEEG can be complemented by other modalities such as concurrent TMS-fMRI. It provides high spatial resolution for noninvasive activity recording of deep brain regions, which allows for investigations in broader neurologically healthy populations. Moreover, while spTMS-evoked responses provide insights into immediate changes in hippocampal activity, they do not reflect longer-term, plasticity-related effects as induced by rTMS ^16^. Thus, combining rTMS with iEEG would be valuable for studying sustained neuromodulatory effects in the hippocampus. However, to our knowledge, no studies to date have implemented this rTMS-iEEG approach in humans.

In the present study, we conducted a series of three experiments using cutting-edge multimodal, concurrent brain stimulation-recording methods (**Fig. 1**). In Experiment 1, we leveraged the concurrent **spTMS-iEEG** (**Fig. 1A–C**) to deliver single pulses of TMS over the parietal cortex in neurosurgical patients and concurrently recorded hippocampal activity changes with iEEG electrodes. The parietal TMS site was either determined by maximal FC to the hippocampus with their individual resting-rest fMRI (hereafter referred to as the “Hippocampal-FC-guided parietal TMS”), or randomly selected or based on maximal FC to other brain regions (referred to as the “Non-Hippocampal-FC-guided parietal TMS”). We tested whether the hippocampal-FC-guided parietal TMS could effectively *target* the hippocampus by evoking significant intracranial activity within this region. Next, in Experiment 2, we employed the concurrent **spTMS-fMRI** (**Fig. 1D–E)** in a healthy cohort to evaluate whether individual differences in intrinsic FC pattern between the parietal TMS site and hippocampus, assessed using resting-state fMRI, could explain the variability in TMS-evoked hippocampal responses. Finally, in Experiment 3, we implemented a novel, first-in-human concurrent **rTMS-iEEG** paradigm (**Fig. 1F–G)** in neurosurgical patients to determine whether the individualized hippocampal-FC-guided parietal rTMS could effectively *modulate* hippocampal activity, specifically by altering theta power, the dominant intrinsic rhythm of the hippocampus ^17, 18^. Together, the results in this study would provide multimodal causal evidence of hippocampal engagement and modulation through individual FC-guided parietal TMS.

**Figure 1.**
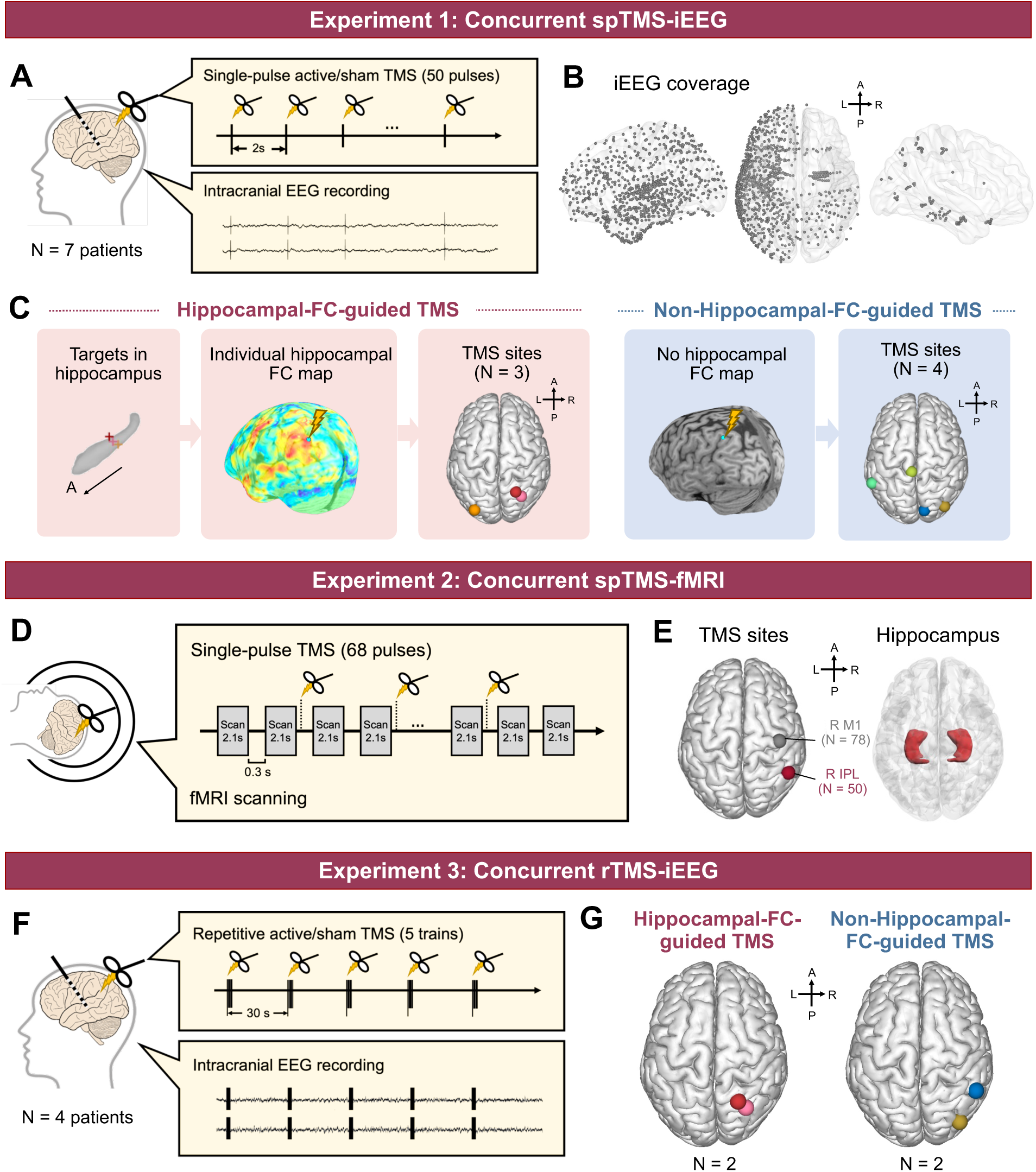
Overview of the multimodal concurrent brain stimulation-recording study design. (**A–C**) In Experiment 1, neurosurgical patients received single-pulse active and sham TMS to their parietal cortex with concurrent intracranial EEG (iEEG) recording in their brain (**A–B**). Their TMS sites were determined by a parietal spot exhibiting individual maximal resting-state FC strength to a hippocampal contact target (“Hippocampal-FC-guided parietal TMS”, **C** left) or a random parietal cortical spot or guided by maximal FC targeting a non-hippocampal contact (“Non-Hippocampal-FC-guided parietal TMS”, **C** right). (**D–E**) In Experiment 2, neurologically healthy participants received single-pulse TMS with concurrent whole-brain fMRI responses recording (**D**). TMS sites included the right inferior parietal lobule (IPL) defined as a group-average frontoparietal network node and the right primary motor cortex (M1) as a control site (**E**). (**F–G**) In Experiment 3, neurosurgical patients with concurrent iEEG recording received either hippocampal-FC-guided or non-hippocampal-FC-guided active and sham repetitive TMS (rTMS) trains to their parietal cortex. See Methods for additional details.

## Results

### Experiment 1 – Concurrent spTMS-iEEG

#### Hippocampal-FC-guided parietal spTMS enhanced intracranial hippocampal responses

For this experiment, 7 neurosurgical patients with iEEG electrodes coverage within the hippocampus received both active and sham spTMS pulses to their individualized hippocampal-FC-guided or non-hippocampal-FC-guided parietal cortex **(Fig. 1A–C**). To evaluate the hippocampal neural responsiveness to parietal TMS, we computed the amplitude of intracranial TMS-evoked potentials (iTEPs) within 25-500 ms window following active and sham TMS in each hippocampal contact for each stimulation strategy. We identified that, 45.0% (9 out of 20) of hippocampal contacts in patients with hippocampal-FC-guided parietal spTMS exhibited strong iTEPs that exceeded 5 standard deviations (SDs) relative to the baseline following the active TMS but not following the sham TMS ^12^. Furthermore, 30.0% (6 out of 20) of hippocampal contacts showed significantly stronger iTEPs following the active TMS as compared to following the sham TMS (*p*_FDR_ < .05) (**Fig. 2A–C**). In contrast, in patients with the non-hippocampal-FC-guided parietal spTMS, only 1 out of 16 hippocampal contacts (6.3%) exhibited strong iTEPs (> 5 SDs) with respect to the baseline, as well as stronger responses to the active than the sham TMS (**Fig. 2F–H**).

**Figure 2.**
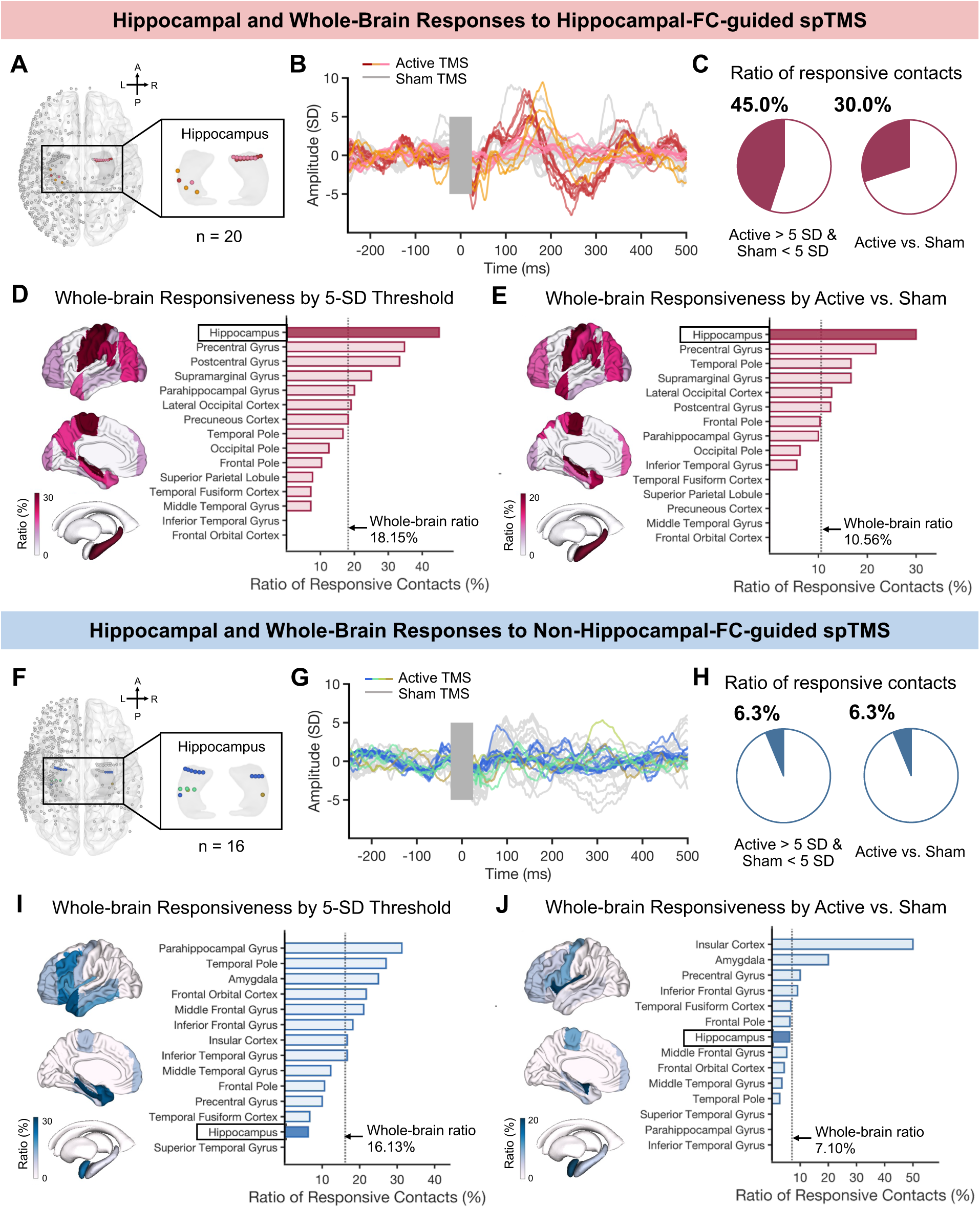
Hippocampal and whole-brain responses to parietal spTMS. (**A**) iEEG electrode contact locations in participants with hippocampal-FC-guided parietal TMS. Hippocampal contacts were highlighted, with different colors representing different participants. (**B**) iTEPs from individual hippocampal contacts following active and sham spTMS. (**C**) Ratio of responsive contacts in the hippocampus based on TMS-evoked potential amplitudes > 5 standard deviations (SD) relative to baseline in the active but not sham condition (left) and active versus sham comparison (right). (**D–E**) The hippocampus was the most responsive region to the hippocampal-FC-guided parietal TMS among all brain areas with sufficient electrode coverage (> 10 contacts across both hemispheres). (**F–J**) Corresponding results from patients with non-hippocampal-FC-guided parietal TMS.

To assess the neurophysiological specificity of hippocampal-FC-guided parietal spTMS on the hippocampus, we expanded the iTEP analyses to all brain regions with sufficient electrode coverage (n > 10 contacts). Across these regions (15 regions covered with n = 303 contacts), the hippocampus emerged as the most responsive region (**Fig. 2D–E)**. The proportion of responsive hippocampal contacts (45.0% and 30.0%) substantially surpassed the whole-brain average: 18.2% based on the 5-SD criterion and 10.6% based on the active versus sham TMS comparison. In contrast, whole-brain analyses in patients with non-hippocampal-FC-guided parietal spTMS (14 regions covered with n = 310 contacts) revealed a relatively low hippocampal response rate, below the whole-brain average rate: 16.1% based on the 5-SD criterion and 7.1% based on the active versus sham TMS comparison (**Fig. 2I–J**). Together, these findings suggest that the individualized hippocampal-FC-guided parietal TMS preferentially engages hippocampal activity.

#### Temporal and spectral profiles of hippocampal responses

After identifying hippocampal contacts responsive to the FC-guided parietal spTMS, we next characterized the temporal and spectral profiles of their neural responses. For temporal dynamics, we performed *post-hoc* point-by-point comparisons of active versus sham iTEP amplitudes across responsive contacts identified with the 5-SD criterion. Three distinct time windows showed significant amplitude differences: 26–49 ms, 108–175 ms, and 223–344 ms (nonparametric cluster-based permutation; cluster-level *t* = –300.5, 409.6, –919.1; *p* = .001, < .001, < .001; 10,000 permutations) (**Fig. 3A**). Moreover, within each significant time window, iTEP amplitudes were substantially greater in responsive contacts compared to non-responsive ones (*t*_(19)_ = –4.12, 4.26, –5.11, *p*_FDR_ < .001) as well as compared to all contacts in patients with non-hippocampal-FC-guided active TMS (*t*_(24)_ = –5.28, 9.53, –6.25, *p*_FDR_ < .001). A summary of iTEP amplitudes relative to the baseline during these time windows for active and sham TMS conditions is provided in **Table S1**.

**Figure 3.**
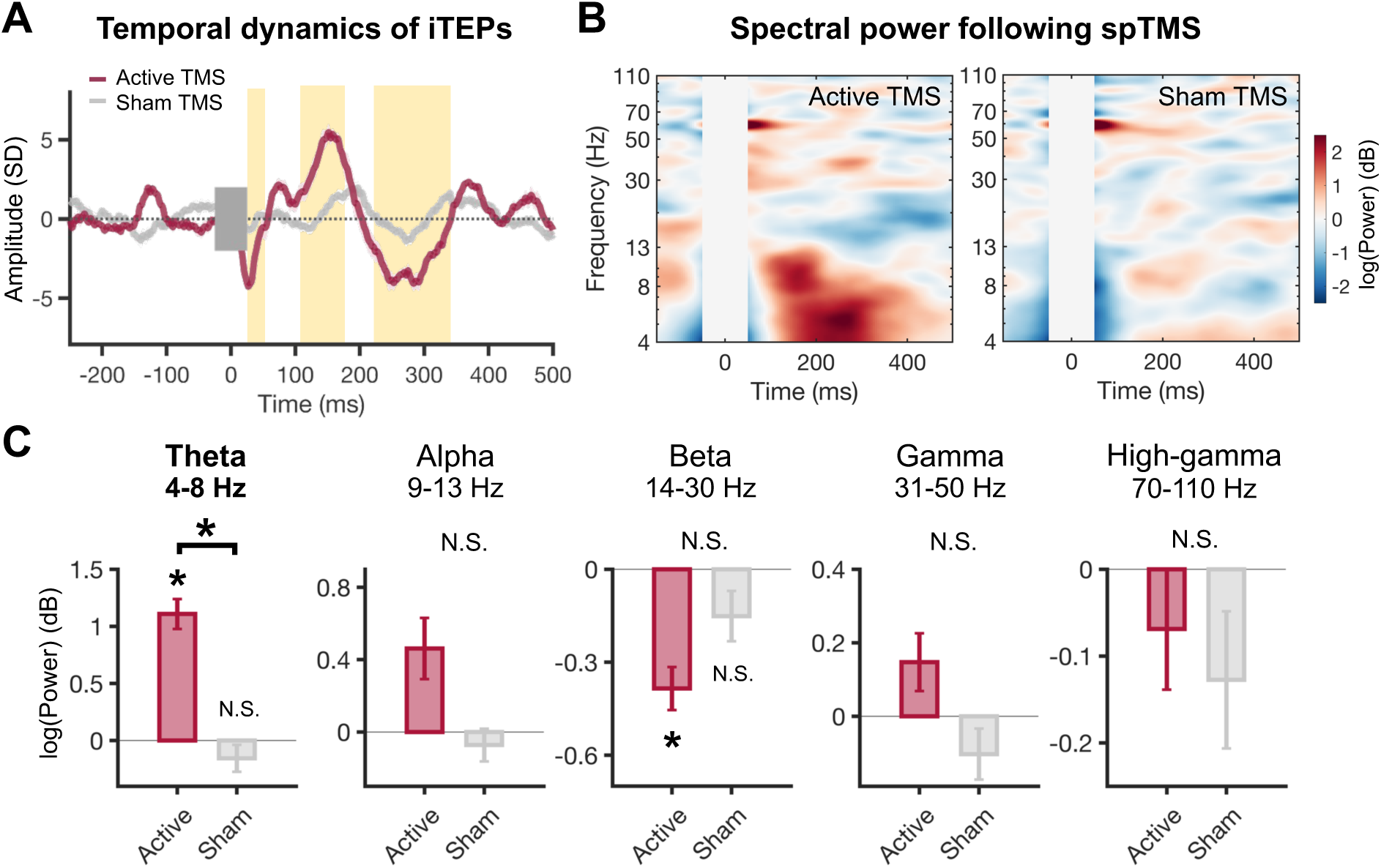
Temporal and spectral profiles of hippocampal responses by hippocampal-FC-guided spTMS. (**A**) Group-level iTEPs across responsive hippocampal contacts. Time windows showing significant active-versus-sham contrast were highlighted. (**B**) Spectral power change relative to the baseline following active (left) and sham (right) spTMS. Artifact period was marked with grey bar. (**C**) Active TMS selectively induced a greater power increase in the theta band compared to that in the sham condition. No significant active-versus-sham contrast was observed in the other frequency bands. Data are presented as mean ± standard error. Asterisks (*) indicate significant results (*p*_FDR_ < .05).

For spectral profiles, we conducted a wavelet-based time-frequency analysis to examine changes in spectral power following parietal spTMS (**Fig. 3B–C**). Power was averaged across a 50–500 ms post-stimulation window within each of five canonical frequency bands: theta (4–8 Hz), alpha (9–13 Hz), beta (14–30 Hz), gamma (31–50 Hz), and high gamma (70–110 Hz). Active TMS induced a significant increase of theta power relative to baseline (*t*_(8)_ = 7.99, *p*_FDR_ < .001), which was also significantly greater than that in the sham condition (*t*_(8)_ = 8.82, *p*_FDR_ < .001). Beta power showed a significant reduction in the active condition (*t*_(8)_ = -5.21, *p*_FDR_ = .004), but this effect did not significantly differ from that in the sham condition (*t*_(8)_ = -1.63, *p*_FDR_ = .23). No significant effect was observed in the other frequency bands (*p*s > .10). These results suggest that hippocampal-FC-guided parietal spTMS specifically influences theta-band oscillations in the hippocampus.

### Experiment 2 – Concurrent spTMS-fMRI

#### Individual FC strength explains parietal spTMS-evoked fMRI responses in the hippocampus

For this experiment, 79 neurological healthy participants received single pulses of TMS inside the MRI scanner while their whole-brain fMRI responses were simultaneously recorded. Of these, 50 participants received TMS to the right inferior parietal lobe (IPL), targeted using an MNI coordinate derived from a group-average parietal node in the fronto-parietal network ^19^. 78 received TMS to the right primary motor cortex (M1), which served as a control site (**Fig. 1E**). Group-level analyses revealed that IPL TMS did not significantly elicit responses in either left or right hippocampus (*t*_(49)_ = –1.67, –0.39; *p* = .10, .69) (**Fig. 4A**) and did not differ from M1 TMS-evoked responses (linear mixed-effect model, *F*_(1, 126)_ = 0.11, 0.13; *p* = .74, .71). These results suggest that a uniform group-based targeting strategy (“one-site-fits-all”) may be insufficient to consistently engage the hippocampus across individuals.

**Figure 4.**
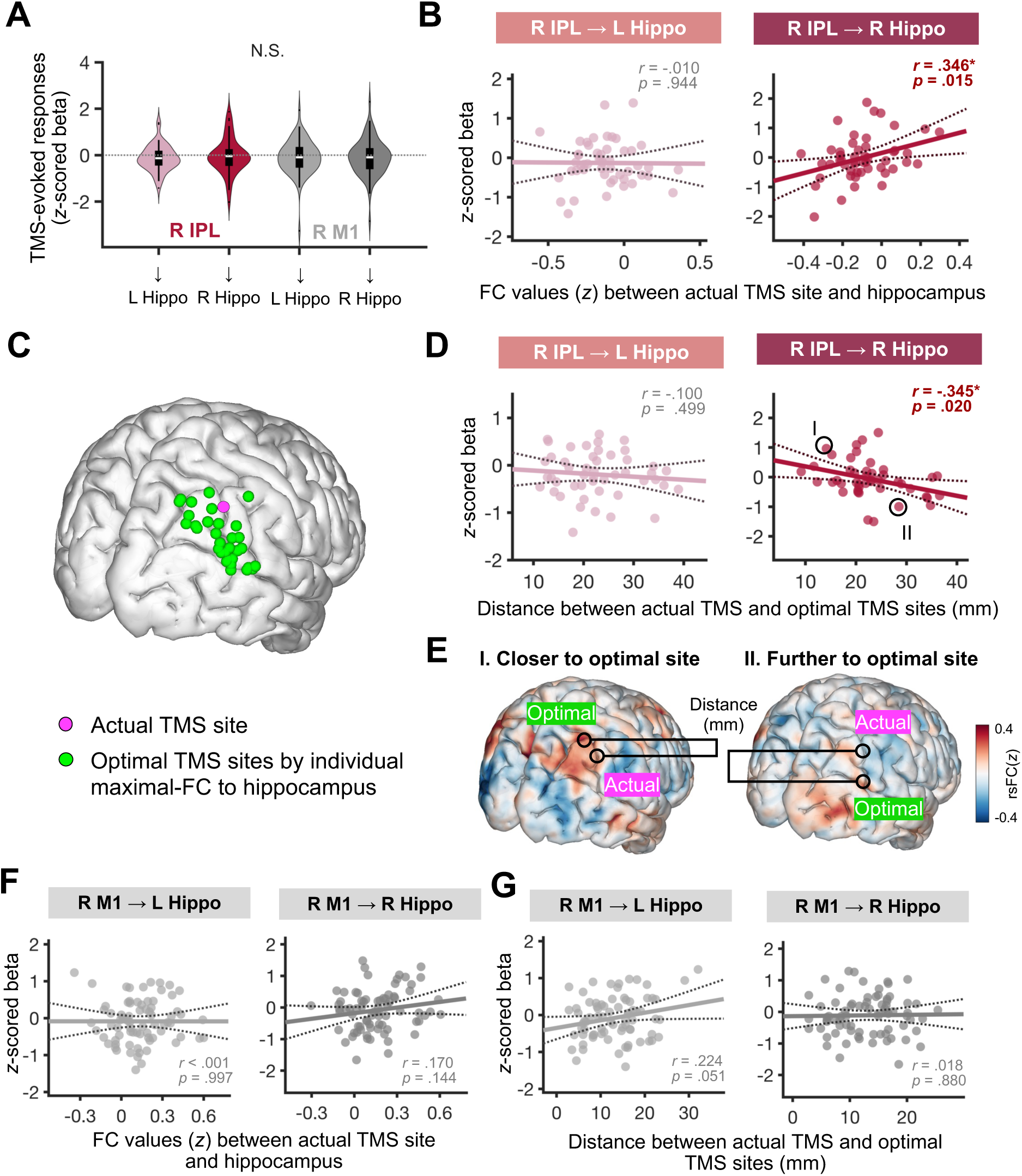
Individual hippocampal-FC explains TMS-evoked hippocampal responses. (**A**) Group-level TMS-evoked responses in the hippocampus following right IPL and right M1 stimulation. (**B**) Right hippocampal responses correlated with FC between the right hippocampus and IPL. No significant correlation was found for the left hippocampus. (**C**) Spatial distribution of each participant’s optimal TMS sites (green dots), defined as the IPL voxels showing maximal FC to the right hippocampus. The group-based actual TMS site was marked with magenta dot. (**D**) Right hippocampal response negatively correlated with the distance between the actual and individualized optimal TMS sites—closer proximity was associated with stronger activation. Two representative participants are outlined and further highlighted in (**E**). (**E**) Two representative participants outlined in (**D**) with the actual TMS site located closer/further to their optimal TMS site defined by their individual hippocampal-FC map. (**F–G**) Control analysis of M1 stimulation. Hippocampal responses were neither correlated with the FC strength between the hippocampus and M1 (**F**), nor with the distance between the actual and optimal sites (**G**).

To investigate the source of this inconsistency, we conducted two follow-up analyses testing whether individual differences in intrinsic hippocampal-FC profile could account for the variability in hippocampal responses to group-targeted IPL TMS. In our first analysis, we assessed each participant’s intrinsic FC between their actual IPL TMS site and hippocampus in each hemisphere using their individual resting-state fMRI and related it to the TMS-evoked hippocampal responses. We identified a significant positive correlation between individual resting-state FC and IPL TMS-evoked responses in the right hippocampus (*Pearson r*_(48)_ = .346, *p* = .015). Specifically, individuals with stronger positive FC between the IPL TMS site and the right hippocampus exhibited greater TMS-evoked responses in the right hippocampus (**Fig. 4B**). No such relationship was observed for the left hippocampus (**Fig. 4B**), or for responses to M1 TMS (*p*s > .09) (**Fig. 4F**).

Previous work has shown that closer proximity between the actual applied and optimal FC-guided TMS sites was related to the clinical response to TMS ^7, 20, 21^, but its relevance to TMS-evoked engagement of a deep target region remains unknown. Therefore, in our second analysis, we investigated whether spatial proximity to a personalized hippocampal-FC-guided parietal stimulation site was associated with the magnitude of TMS-evoked hippocampal responses. As in Experiment 1, we identified the personalized stimulation site as the parietal cortical spot showing maximal positive resting-state FC to the hippocampus (**Fig. 4C**) and quantified the spatial proximity as the Euclidean distance between the actual and personalized stimulation sites. We identified a significant negative correlation between this distance and TMS-evoked responses in the right hippocampus (*Pearson r*_(44)_ = –.345, *p* = .020), indicating that closer proximity to the personalized FC-guided parietal site was associated with greater right hippocampal responses (**Fig. 4D**). No such a relationship was observed in the left hippocampus, or for M1 TMS (*p*s > .05) (**Fig 4G**). Overall, these findings underscore the value of using individualized FC-guided targeting to optimize parietal TMS for effective hippocampal engagement.

### Experiment 3 – Concurrent rTMS-iEEG

#### Hippocampal-FC-guided parietal rTMS reduced hippocampal theta-band power

To evaluate the modulation effects of different stimulation strategies on hippocampal activity, we applied 5 active and 5 sham rTMS trains (10 Hz or 20 Hz) to hippocampal-FC-guided or non-hippocampal-FC-guided parietal cortex in 4 neurosurgical patients (**Fig. 1F**). TMS-induced changes of hippocampal spectral power were assessed relative to the baseline under each stimulation strategy (**Fig. 5A**) and averaged over all five 25 secs post-rTMS time windows within each of five canonical frequency bands (**Fig. 5B**). In patients with hippocampal-FC-guided parietal rTMS, active stimulation significantly reduced theta power across hippocampal contacts (*n* = 18; *t*_(17)_ = –10.74, *p*_FDR_ < .001), with no effect for sham stimulation (*t*_(17)_ = –0.41, *p*_FDR_ = .78). Moreover, theta power suppression was significantly greater following active than sham stimulation (*t*_(17)_ = –10.56, *p*_FDR_ < .001) (**Fig. 5B**), with no significant effects in other frequency bands for either stimulation condition or between conditions (*p*s > .10) (**Fig. 5B**). Additionally, we observed a prominent cumulative modulation effect: theta suppression following the final train was significantly greater than that following the 1^st^, 3^rd^, and 4^th^ trains (*t*_(17)_ = 6.15, 2.94, 4.91; *p*_FDR_ < .001, = .030, < .001), and showed a trend of stronger level than the 2^nd^ train (*t*_(17)_ = 1.94, *p*_FDR_ = .14) (**Fig. 5C**). In contrast, non-hippocampal-FC-guided parietal rTMS did not induce significant theta power change (*n* = 6; *t*_(5)_ = –2.53, *p*_FDR_ = .26) (**Fig. 5D–E**). Direct comparison confirmed a significantly stronger theta suppression with hippocampal-FC-guided than non-hippocampal-FC-guided parietal active rTMS (*t*_(22)_ = –3.01, *p*_FDR_ = .048) (**Fig. 5F**), with no significant group differences in other frequency bands (*p*s > .16).

**Figure 5.**
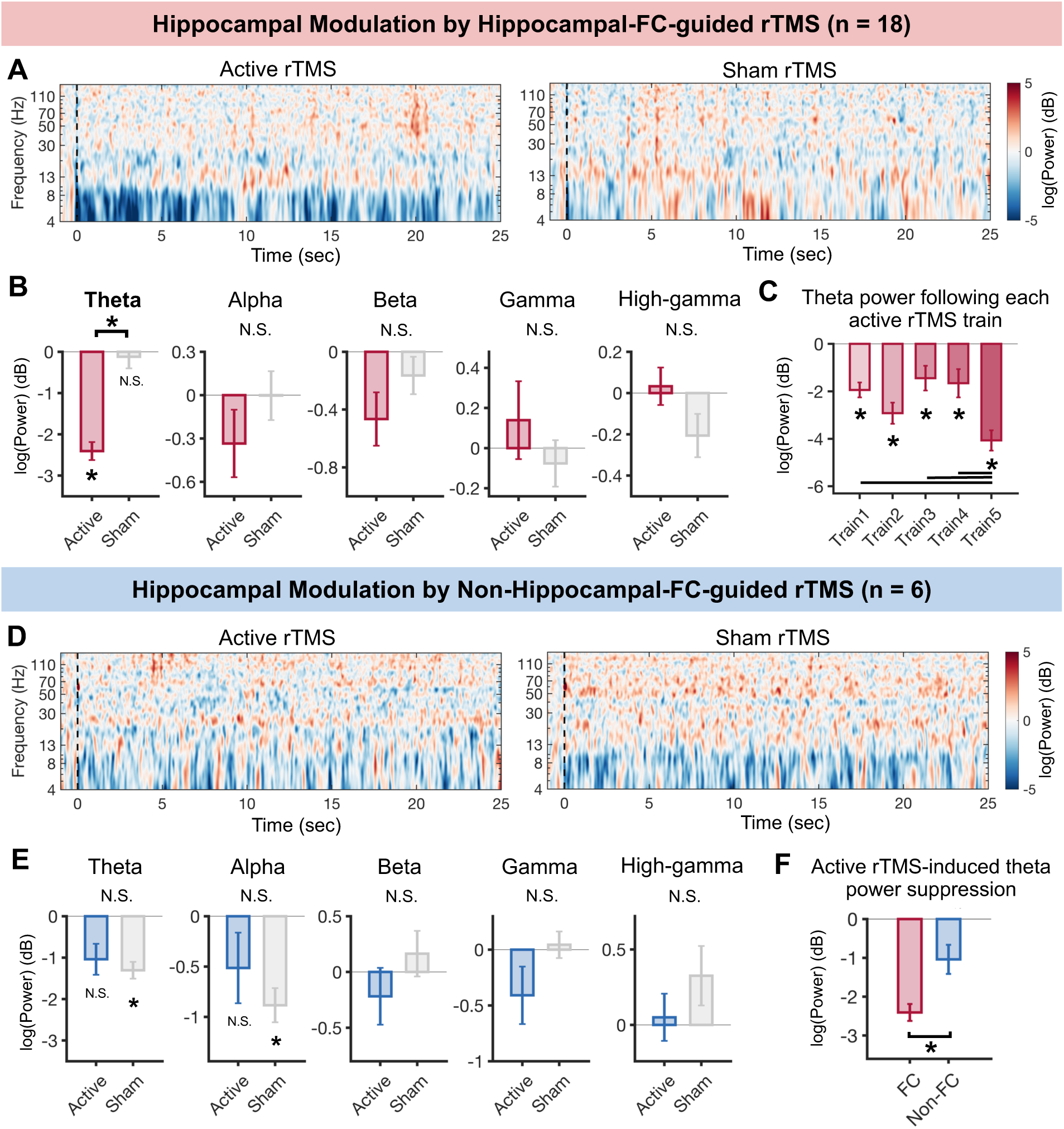
Hippocampal modulation by hippocampal-FC-guided parietal rTMS. (**A**) Spectral power in the hippocampus following hippocampal-FC-guided rTMS. (**B**) Hippocampal-FC-guided active rTMS significantly suppressed theta power, with no effects in other frequency bands. Data are represented as mean ± standard error. (**C**) Theta power suppression following each active rTMS train, with the strongest effect following the final train. Asterisks (*) indicate significant power relative to zero (baseline) (*p*_FDR_ < .05). Horizontal bars indicate significant differences between trains (*p*_FDR_ < .05). (**D–E**) Spectral power following non-hippocampal-FC-guided rTMS. No significant effects were observed in the active condition. (**F**) Theta suppression was significantly greater following hippocampal-FC-guided rTMS than non-hippocampal-FC-guided rTMS.

To evaluate the persistence of rTMS-induced theta modulation, we computed theta power suppression in successive 1-sec windows post-stimulations. Significant theta suppression lasted for over 20 secs after each train (**Fig. S3**). To further evaluate the modulation specificity of hippocampal-FC-guided TMS on the hippocampus, we examined theta power changes in the parahippocampal gyrus and amygdala (**Fig. S4**). No significant theta power change was observed in these regions (parahippocampal gyrus: *t*_(6)_ = –3.23, *p*_FDR_ = .07; amygdala: *t*_(8)_ = –1.44, *p*_FDR_

= .33). The theta power suppression was significantly greater in the hippocampus compared to the parahippocampal gyrus (*t*_(23)_ = –2.11, *p*_FDR_ = .045) and the amygdala: *t*_(25)_ = –4.43, *p*_FDR_ = .004) (**Fig. S4C**). Overall, these results suggest that hippocampal-FC-guided rTMS induced sustained and selective hippocampal theta activity.

## Discussion

We investigated whether hippocampal-FC-guided parietal TMS can effectively engage and modulate the human hippocampus through a series of multimodal concurrent brain stimulation-recording experiments. In particular, the concurrent TMS-iEEG provides powerful, direct measurements and compelling evidence from neurosurgical patients that hippocampal-FC-guided parietal TMS enhanced both hippocampal activity and plasticity. Complementary noninvasive evidence integrated from concurrent TMS-fMRI and individual resting-state fMRI measurements in neurologically healthy participants further showed intrinsic FC between the stimulation site and hippocampus accounted for variability in TMS-evoked hippocampus response. These findings demonstrated the effectiveness of an individualized, hippocampal-FC-guided stimulation strategy for precise targeting and modulation of the hippocampus.

Hippocampal-FC-guided parietal TMS has previously been shown to change hippocampal network connectivity ^5, 6^, i.e., communication between brain regions. However, such findings do not establish a causal relationship between regions. According to the theory of “coherence through communication” ^22^, change in inter-regional connectivity could be epiphenomenal, resulting from local alterations in synaptic coupling and oscillatory power at the stimulated region ^10^, rather than reflecting true downstream effects at the target region. Our findings have two key advantages that address the limitations of prior work on the modulation of hippocampal network connectivity. First, we provide direct causal evidence that hippocampal-FC-guided parietal TMS can specifically engage the hippocampus. Nearly half of hippocampal contacts exhibited significant iTEPs relative to the baseline within 500 ms following single pulses of active TMS to the hippocampal-FC-guided parietal cortex, whereas no such effects were observed following sham TMS. Furthermore, one-third of hippocampal contacts also showed significantly stronger iTEPs in the active versus sham TMS condition. This proportion was substantially higher than the 6.3% observed with the hippocampal-FC-guided parietal TMS, and the ∼6% reported in our previous study using non-FC-guided DLPFC TMS ^14^. We further demonstrated that the hippocampus showed the highest percentage of responsive contacts among all recorded brain areas, reinforcing the utility of targeted FC-guided cortical TMS for enhancing selective engagement of deep brain regions like the hippocampus.

Second, we demonstrated enhanced hippocampal activities independent of any task. While some may view this as a limitation of this study in understanding how stimulation influences memory function, task-free stimulation provides a clearer window into the direct neural effects of TMS. Specifically, it helps disentangle downstream effects on the target region from confounding activity related to task-dependent behaviors. However, future studies should still explore whether and how FC-guided TMS can directly influence task-related hippocampal activity and, in turn, impact ongoing hippocampus-dependent memory behavior.

Interestingly, we observed an early iTEP component (emerging within 30 ms), as well as two later components (around 150 ms and 250 ms), in responsive hippocampal contacts but not in nonresponsive contacts following hippocampal-FC-guided parietal TMS. These temporal patterns suggest that the propagation of hippocampal-FC-guided parietal TMS to the hippocampus may involve either monosynaptic or polysynaptic pathways. Specifically, the early component likely reflects a direct axonal, likely monosynaptic connection (^23^ but see also ^24^) from the parietal cortex to the hippocampus, while the later components may reflect indirect polysynaptic transmission through a cascade of intermediate regions ^23^, such as the pulvinar thalamus, perirhinal cortex, parahippocampal cortex, and retrosplenial cortex that have been involved in hippocampal network connectivity ^9^. Supporting this interpretation, spectral analysis revealed that parietal spTMS elicited a selective increase of hippocampal theta oscillations, which are hallmark electrophysiological features of both rodent and human hippocampus and support memory functions by regulating inter-regional communication within the hippocampal network ^17, 18^. Together, these findings from the concurrent spTMS-iEEG approach provide direct evidence that hippocampal-FC-guided parietal TMS causally engages the hippocampus, highlighting its potential to modulate hippocampal activity and associated functions.

Importantly, our findings in neurologically healthy participants with concurrent TMS-fMRI provided solid neural evidence supporting a personalized, FC-guided TMS targeting strategy. Historically, the “one-site-fits-all” approach has dominated TMS targeting due to clinical practical constraints, which predominantly focused on the DLPFC-sgACC circuit. However, accumulating evidence now points to inter-individual variability in both brain connectivity ^25^ and treatment outcomes ^26^. For example, prior work has shown that the strength of individual FC between the DLPFC and the sgACC predicts antidepressant response to DLPFC TMS ^27, 28^. Compared to traditional anatomical targeting methods, the individual FC-guided TMS strategy can better optimize (anti)correlations between the stimulation site and downstream targets ^29^, thereby enhancing the efficacy of the TMS. Supporting this, stimulation sites with closer spatial proximity to an individual’s peak sgACC anti-correlated DLPFC spot are associated with greater clinical improvement ^20, 21^. In line with this literature, we did not observe significant TMS-evoked responses in the hippocampus using a group-level coordinate for parietal stimulation. We observed substantial individual variation in the TMS-evoked hippocampal response, which was positively associated with the individual resting-state FC strength between the actual parietal stimulation site and hippocampus. Moreover, the spatial proximity between the actual TMS site and the optimal site, i.e., the individual’s peak parietal spot showing maximal positive FC to the hippocampus, predicts the magnitude of TMS-evoked hippocampal responses. These findings not only validate our observations in clinical populations (e.g., neurosurgical patients) but also demonstrate that intrinsic FC can be leveraged both to optimize TMS target engagement and as a predictive biomarker of downstream neural activation ^11^. Thus, our work broadens the application of FC-guided TMS from frontal-limbic to posterior-medial temporal circuits, paving the way for targeted neuromodulation of memory-related networks.

Our concurrent rTMS-iEEG findings from Experiment 3 further corroborate the potential of the hippocampal-FC-guided parietal TMS in its capacity to modulate hippocampal oscillations. rTMS is assumed to produce lasting neural effects via synaptic or non-synaptic plasticity mechanisms ^16^. We found that both 10 and 20 Hz active rTMS delivered to an individually defined hippocampus-FC-guided parietal site of neurosurgical patients led to a sustained theta-band oscillatory change in the hippocampus, lasting over 20 secs post-stimulation. This plasticity effect was not seen in other frequency bands, or in sham rTMS condition, or non-hippocampus-FC-guided rTMS condition, or in neighboring regions. These findings extended the specificity of *transient* hippocampal engagement by hippocampal-FC-guided parietal spTMS observed in Experiment 1. Moreover, our study provides the first-in-human evidence of rTMS-induced plasticity effects in downstream targets with direct recording with iEEG. Specifically, we observed a suppression of theta power in the hippocampus, rather than a typical facilitation of neural activity following high-frequency rTMS (> 5 Hz), which was mainly examined and restricted in the local stimulation region ^16^. This suppression effect can be attributable to the timing of stimulation relative to the endogenous oscillatory phase. Intracranial evidence from both rodents ^31^ and humans ^32^ has shown that stimulation at different hippocampal theta phases, e.g., peak versus trough, induced different plasticity effects of facilitation or inhibition in the hippocampus. Supporting this, a recent close-loop neuromodulation work ^33^ reported reductions in hippocampal theta power following repetitive, phase-blind cortical stimulation, which is in line with our result with a non-phase-locked rTMS protocol. Future FC-guided TMS studies that incorporate phase-locking to hippocampal theta rhythms hold promise for more precise modulation of plasticity processes, enabling selective facilitation or inhibition effects. Nevertheless, our findings of selective modulation of hippocampal theta oscillations by parietal rTMS align with a recent study that used invasive parietal stimulation ^34^, thus extending this effect to the noninvasive stimulation approach. These results together position hippocampal-FC-guided parietal TMS as a promising method to modulate hippocampus-dependent functions through noninvasive and selective perturbation of ongoing hippocampal theta oscillations.

There are several limitations to this work. First, the sample sizes of the neurosurgical patients in our iEEG experiments are limited due to practical constraints inherent to enrolling a specific population requiring epilepsy surgery. Future efforts involving multi-site collaborations may help overcome this limitation. Second, while our findings suggest that hippocampal modulation induced by hippocampal-FC guided rTMS can persist for tens of seconds, it remains unknown whether these effects extend beyond several minutes or translate into longer-term plasticity. Future work should incorporate longer post-stimulation monitoring windows to better evaluate the durability and stability of these plasticity effects. Finally, this work employed conventional high stimulation frequency (10 Hz / 20 Hz) as a preliminary exploration. Future research could examine the feasibility of tailoring TMS frequency to individuals’ intrinsic oscillatory profiles, in conjunction with the FC-guided TMS targeting. Aligning stimulation frequency with the endogenous oscillatory dynamics of a targeted region or network shows potential of enhancing the propagation and efficacy of TMS-induced effects ^35^.

In conclusion, our study provides direct, multimodal evidence that individualized, FC-guided parietal TMS can causally engage and modulate human hippocampal activity. These findings lay critical groundwork for developing precision neuromodulation strategies aimed at enhancing hippocampus-related cognitive functions, with promising applications for treating deficits associated with dementia, memory impairment, and affective disorders. More broadly, our results underscore the value of an individualized, connectivity-informed approach to noninvasive brain stimulation, with far-reaching implications for both basic neuroscience and clinical practice.

## Acknowledgments

JJ is supported by the National Institutes of Health (R01MH136197) and the Brain and Behavior Research Foundation Young Investigator grant (29441). NTT is supported by grants from the National Institute of Mental Health (1K23MH125145 and R01MH125160), Magnus Medical, Inc., and the Brain and Behavior Research Foundation (31275). AE is supported by the National Institutes of Health (R01 MH103324). ADB is supported by the National Institutes of Health (R21MH120441 and R01NS114405) and Roy J. Carver Trust. Other funding includes NIMH R01MH132074 (NTT and ADB). This work was conducted, in part, on an MRI instrument funded by 1S10OD025025-01.

We would like to acknowledge the patients and families who graciously agreed to participate in this research, as well as research members in the Human Brain Research Laboratory who assisted with data collection and provided feedback on the project including Ariane Rhone, Haiming Chen, Ben Pace, Brandt Uitermarkt, Kirill Nourski, Joel Berger, Hiroyuki Oya, and Chris Garcia.

## Author contributions

Conceptualization, ZL, ADB, JJ; Writing – original draft, ZL, JJ; Writing – review and editing, all authors; Investigation, ZL, NTT, JB, MT, ADB, JJ; Formal analysis, ZL, JJ; Methodology, ZL, JB, XL, KW, ZC, JJ; Funding acquisition, resources, and supervision, NTT, AE, MAH, ADB, JJ

## Code Availability

All codes for preprocessing are provided at https://github.com/JingjiangLab/Parietal-Hippocampus.git

## Supplemental information

Document S1: Figure S1-S4, and Table S1-S4.

## Methods

### 1. Concurrent spTMS-iEEG Experiment

#### Participants

In total, 11 adult patients with medically intractable epilepsy enrolled in this study and received TMS over parietal cortex while concurrently recording brain signals with iEEG. These patients were admitted to the University of Iowa Hospitals and Clinics for 14 days of monitoring with intracranial electrode implantation to localize their seizure focus. Four patients were later excluded due to either 1) a lack of sham condition (1 excluded), 2) excessive background noise with use of a different iEEG amplifier (1 excluded), 3) hippocampus electrodes at seizure focus (1 excluded), or 4) frequent seizures on the experiment day, including during data collection (1 excluded). The final 7 patients (2 females, age 37.4 ± 15.1) included for main analyses passed stringent visual and quantitative data quality control procedures. The demographics of these patients are shown in **Table S2**. All experimental procedures were approved by the University of Iowa Institutional Review Board (IRB) and were under ethical principles in the Declaration of Helsinki. Written informed consent was obtained from all participants.

#### Experimental design

Prior to intracranial electrode implantation, patients underwent an MRI and resting-state functional MRI (rs-fMRI) scan. The rs-fMRI data were used to identify personalized stimulation targets within the parietal area that showed maximal functional connectivity to the hippocampus. These personalized hippocampal FC-guided targets were used for the concurrent TMS-iEEG experiment. The day following electrode implantation, patients underwent a second MRI scan and thin-slice volumetric computerized tomography (CT) scan for clinical purposes to identify the location of the electrode contacts. The TMS-iEEG experiment was conducted after seizures were localized, typically 1-2 days before electrode explantation and 24 hours after restart of anti-seizure medications. The safety and feasibility of this experimental method have been demonstrated by our prior work ^14^. An overview of the experimental design is illustrated in **Fig. 1A-C**.

#### Pre- and post-implantation imaging scans

Pre-implantation structural MRI was acquired using a 3T GE Discovery 750 W scanner with a 32-channel head coil. A 3D FSPGR BRAVO sequence was used: TR = 8.49 ms, TE = 3.28 ms, TI = 450 ms, FOV = 25.6 cm, Flip angle = 12°, voxel size = 1.0 × 1.0 × 0.8 mm. Twenty-five minutes of rs-fMRI scan was acquired for 6/7 patients through five 5-minute gradient-echo EPI runs (650 volumes): TR = 2260 ms, TE = 30 ms, FOV = 22.0 cm, Flip angle = 53°, 60 slices (2.5 mm thick, voxel size = 3.44 × 3.44 × 4 mm) covering the whole brain. Post-implantation MRI was acquired using a 3T Siemens Skyra scanner with a head transmit-receive coil. An MPRAGE sequence was used: TR = 1970 ms, TE = 3.44 ms, TI = 1000 ms, Flip angle = 10°, FOV = 25 cm, voxel size = 1.0 × 1.0 × 0.8 mm. The CT acquisition voxel size was 0.47 × 0.47 × 1.0 mm.

#### MRI processing

Standard MRI preprocessing was performed for the pre-implantation scans using the fMRIPrep pipeline ^36^ as follows. The T1-weighted (T1w) image was processed with ANTs ^37^ for intensity nonuniformity correction, skull-stripping, and spatial normalization to the MNI template through nonlinear registration. Cerebrospinal fluid (CSF), white-matter (WM) and gray-matter (GM) were segmented on the brain-extracted T1w using FAST (FSL). For each of the rs-fMRI run data, preprocessing steps included removal of the first 6 volumes, head-motion estimation using MCFLIRT (FSL), co-registration to the T1w reference using boundary-based registration with FLIRT (FSL), distortion correction using a blip-up/blip-down “phase difference” fieldmap acquisition, and resampling onto the standard MNI space. Confounding variables such as framewise displacement (FD) ^38^, DVARS and three global signals from the CSF, the WM, and the whole brain were also estimated.

The following post-processing steps were performed using XCP-D ^39, 40^. First, 36 confounding regressors were selected from fMRIPrep outputs. They included 6 motion parameters, 3 mean signals from the CSF, the WM, and the whole brain, temporal derivatives of these 9 parameters, and the quadratic expansion of both the 9 parameters and their temporal derivatives. Next, high-motion outlier volumes were identified with FD value > 0.5 mm and used for data interpolation. The interpolated data and confounds are then detrended and mean-centered, followed by 0.008-0.08 Hz bandpass filtering. The filtered confounds were then regressed from the filtered BOLD data. Finally, the resulting data were censored (high motion volumes removal) and smoothed with a 6-mm FWHM Gaussian kernel.

#### Intracranial electrodes implantation and localization

Patients were implanted with depth electrodes (Stereo-electroencephalography, sEEG) and/or subdural grid/strip arrays (electrocorticography, ECoG) (Ad-Tech Medical, Racine, WI) (**Table S3**). Each depth electrode shaft consisted of 4 to 10 platinum macro-electrode contacts (1.3 mm diameter, 1.6 mm length, 5–10 mm inter-electrode distance). Subdural arrays were flexible silicon sheets embedded with platinum-iridium disc-shaped contacts (2.3 mm diameter, 5–10 mm inter-contact distance).

To precisely determine locations of intracranial electrode contacts for each patient, the post-implantation MRI and CT imaging was linearly co-registered to pre-implantation MR scans using FLIRT (FSL). Images were carefully corrected for displacement and distortion caused by electrodes implantation using non-linear 3D thin-plate spline warping with 50-100 manually selected control points. Coordinates of each electrode contact identified on the native space of patients’ post-implantation MRI and CT scans were thus projected onto the pre-implantation MRI space and then normalized to standard MNI (ICBM152 template) space using ANTS ^37^. Contacts with MNI coordinates falling within the hippocampus mask, as defined by the probabilistic maps of the Harvard Oxford Structural Atlases at a 25% threshold in the FSL^41^, were initially identified as hippocampal contacts of interest. The location of each selected contact was visually reviewed in each participant’s native space to further confirm its location.

#### Hippocampal FC-guided TMS site definition

Patients were randomly assigned into two groups based on different TMS targeting strategies. In the experimental group of 3 patients, we defined personalized parietal stimulation targets that showed maximal rs-fMRI connectivity to a hippocampal seed for each patient, as done in previous work ^5, 6^. Specifically, the contacts within the left hippocampus were first identified and a 4-mm radius sphere was placed at the contact nearest to MNI [−29, −25, −13], which served as the hippocampal seed for each patient. This coordinate was previously used in previous literature ^6^ as a representative voxel of the hippocampal regions. Next, this hippocampal seed was used to generate an individual FC map of hippocampal resting-state functional connectivity (FC). The FC was computed based on the Fisher’s *z*-transformed Pearson correlation coefficients between the BOLD times series of this seed and every voxel of the brain from the post-processed rs-fMRI data in native space. A cortically accessible stimulation location was selected as the voxel showing the strongest positive hippocampal connectivity within the left parietal cortex, or right parietal cortex if the left parietal was not accessible for TMS because of electrode anchors impeding access. This strategy resulted in 1 patient receiving hippocampus-FC guided TMS in the left parietal cortex and 2 patients receiving that in the right parietal cortex (**Fig. 1C**).

For the control group, 4 patients received TMS either at a random parietal cortical spot (2 patients) or guided by individual FC targeting a non-hippocampal target within the temporal lobe (1 patient targeting precuneus, 1 patient targeting parahippocampal cortex) (**Fig. 1C**). Of these, 2 patients received TMS in the left parietal cortex and 2 patients in the right. The MNI coordinates of all parietal stimulation sites are provided in **Table S3**.

#### Concurrent TMS and iEEG recording

A Cool-B65 Active/Placebo (A/P) liquid-cooled butterfly coil that connected to a MagVita X100 230V stimulator (Magventure, Alpharetta, GA, USA) was used for delivering TMS pulses. Brainsight Neuronavigation System (Rogue Research, Montreal, Quebec, Cannada) with frameless stereotaxy was used to guide the stimulation to targets identified on the pre-implantation structural MRI images. Prior to the experiment, motor threshold (MT) was defined for each patient as the stimulation intensity that induced visually observable hand movements in at least 3 of 5 consecutive trials by stimulating the hand knob of motor area.

In the TMS-iEEG experiment, 50 single pulses of TMS at 0.5 Hz were delivered either to the hippocampal FC-guided or non-hippocampal-FC-guided parietal location defined above. For the hippocampal FC-guided parietal targeting, the individual hippocampal FC map was loaded into Brainsight to pinpoint the spot with strongest positive hippocampal connectivity within the parietal cortex for stimulation. Stimulation intensity was applied at 120% MT or 100% MT if 120% was not tolerated due to discomfort). As a control condition, 50 sham pulses were also delivered to the same target with the coil flipped 180°, directed away from the head. Earplugs were used in both conditions to reduce the influence of auditory confounds. During both conditions, iEEG data acquisitions were simultaneously made by a multichannel data acquisition system (ATLAS, Neuralynx, Tucson, AZ) with a reference contact placed in patients’ subgaleal space. The sampling rate was 8 kHz or 4 kHz with 1–2000 Hz acquisition filters (−6 dB, 256 tap length).

#### iEEG preprocessing

Preprocessing was performed using a customized MATLAB pipeline based on the FieldTrip toolbox ^42^, following procedures consistent with our prior work ^14^. First, contacts located within the seizure onset zone or exhibiting prominent epileptic artifacts were excluded from analysis. For the remaining contacts, the TMS artifact window (−10 to 25 ms relative to TMS onset) was removed, resulting in segments of iEEG signal from 26 to 1990 ms post-stimulation onset. Signals were then filtered using a 3rd-order Butterworth notch filter to remove line noise at 60 Hz and its harmonics (57–63 Hz and seven harmonics), followed by a 2nd-order Butterworth bandpass filter (2–35 Hz, 6 dB cutoff). The missing artifact period (−10 to 25 ms) was interpolated using segments both preceding and following the artifact window. The segments were weighted with tapering functions and then summed to generate a replacement that preserved signal stationarity^43^. After interpolation, the data were downsampled to 1000 Hz and epoched from −250 to 500 ms relative to TMS onset. Then, residual decay artifacts were removed using the Adaptive Detrend Algorithm (ADA ^44^). Finally, trials contaminated by non-physiological artifacts (e.g., cable motion artifact, long decay artifact from amplifier saturation) or potential interictal spikes were discarded. Specifically, trials were rejected if the signal exceeded 10 times the baseline (−250 to −50 ms) standard deviations (SDs) or 200 µV at 26 ms, or 50 times the baseline SD or 300 µV throughout the entire epoch. Contacts with fewer than 50% trials remained were discarded from further analysis.

#### Quantification and Significance Testing of Intracranial TMS-Evoked Potentials (iTEPs)

After preprocessing, iTEPs were computed by averaging the remaining individual trials for each contact. The resultant iTEPs were then *z*-score normalized with respect to their baseline SDs. To quantify the evoked responses in the hippocampus, we first analyze the responsiveness of each hippocampal contact. Two criteria were employed: 1) Threshold criterion: Contacts were classified as responsive if their iTEPs between 26 ms and 500 ms exceeded 5 baseline SDs under the active TMS condition, but not under the sham condition; 2) Contrast criterion: A two-sample *t*-test (two-sided) was performed to compare active and sham iTEP amplitudes point-by-point across all trials over 26−500ms. For each trial, the baseline amplitude was subtracted from the iTEPs. The resulting *p*-values were corrected using the false discovery rate (FDR). Contacts exhibiting significant active versus sham differences were identified as responsive. Then, for each criterion, the number of responsive contacts was divided by the total hippocampal contacts, yielding the percentage of responsive contacts in the hippocampus.

To further identify the time window(s) of significant iTEPs, we employed a nonparametric cluster-based permutation test ^45^ comparing active and sham TMS conditions across the responsive hippocampal contacts. Paired *t*-tests (two-sided) were first conducted at each post-stimulation time point between 25 and 500 ms. Temporally adjacent time points exceeding an uncorrected alpha level of 0.05 and spanning more than 10 ms were merged into clusters. For each cluster, a cluster-level *t*-statistic was calculated by summing the *t*-statistic of each time point within the cluster. To assess the statistical significance of these clusters, we generated a null distribution of cluster-level *t*-statistics by randomly shuffling trials for 10,000 permutations. Specifically, trial labels (active versus sham) were randomly shuffled for each contact. The iTEPs were recomputed and then compared across conditions. In each permutation, only the maximal cluster-level statistic from each permutation was saved to generate the null distribution. Observed clusters were considered significant if their *p*-values, derived from this null distribution, were below 0.05.

#### Time-frequency analysis of iTEPs

To further examine which specific frequency ranges of hippocampal neural activity were influenced by hippocampal-FC-guided parietal spTMS, we performed a time-frequency analysis on the significant hippocampal iTEPs. Specifically, we applied time-frequency decomposition to trial-level responses using Morlet wavelets (4–110 Hz), implemented with custom MATLAB scripts and the EEGLAB toolbox ^46^. The wavelet cycle parameters were set to [2, 0.5], allowing more cycles for higher frequencies to ensure comparable temporal resolution of the estimation across the frequency spectrum. For each contact, trial-averaged spectral power was computed and baseline-corrected by subtracting the mean within the temporal window [–500, –50 ms]. The resulting power was then averaged within the window of 50 to 500 ms post-stimulation and within five canonical frequency bands: theta (4–8 Hz), alpha (9–13 Hz), beta (14–30 Hz), gamma (31–50 Hz), and high gamma (70–110 Hz). To assess frequency-specific effects, one-sample *t*-tests (two-sided) were first conducted separately for the active and sham conditions across hippocampal contacts. Subsequently, the power values were compared across TMS conditions (active versus sham) and across TMS strategies (Hippocampal-FC-guided versus non-hippocampal-FC-guided), thereby identifying frequency-specific neural effects induced by hippocampal-FC-guided active spTMS.

### 2. Concurrent TMS-fMRI Experiment

#### Participants

In this study, 79 healthy right-handed adult participants were recruited via advertisements. Trained doctoral-level clinicians administered clinical interviews to determine the absence of current and past psychiatric or neurological disorders, claustrophobia, and regular use of psychiatric medications. 50 (30 females, age 31.8 ± 11.3) and 78 (48 females, age 31.4 ± 10.3) participants were included in the final main analyses for the effects of stimulating parietal and M1 respectively. The relatively lower number of participants for parietal stimulation was due to this site being less accessibility by TMS within the MRI scanner head coil for certain participants with larger head sizes. Demographics of participants for each stimulation are shown **Table S4**. All participants signed informed consent. All experimental procedures were approved by the Stanford IRB and were conducted under ethical principles in the Declaration of Helsinki.

#### Experimental design

Participants were invited for two scan visits on two separate days. During the baseline scan day, high-resolution structural MRI and rs-fMRI images were collected. The rs-fMRI data were used to retrospectively evaluate the hypothesis that the TMS-evoked hippocampal response assessed via the TMS-fMRI experiment could be explained by the parietal-hippocampus FC. During the TMS-fMRI scan day, TMS was delivered over the participant’s parietal area and M1 in the MRI scanner in concurrent with fMRI data acquisition to assess single-pulse TMS-evoked response in the hippocampus. An overview of the experimental design is illustrated in **Fig. 1D-E**.

#### Visit 1: Magnetic resonance imaging

We collected structural MRI and rs-fMRI of each participant using a 3T GE 750 scanner with an 8-channel head coil during their baseline scan day. The structural MRI was acquired using high resolution T1 weighted (T1w) (3D inversion SPGR) sequence: TR = 8.6 ms, TE = 3.4 ms, TI= 450 ms, flip angle = 15°, FOV = 22 cm, 184 slices, matrix 256 × 256, 1.0 × 0.9 × 0.9 mm acquisition resolution. The rs-fMRI run was acquired using a T2-weighted (T2w) gradient-echo spiral in/out pulse sequence: TR = 2000 ms, TE = 30 ms, flip angle = 80°, FOV = 22 cm, pixel size = 3.4 mm, 31 axial slices (4 mm thick) covered the whole brain, slice spacing 0.9 mm, matrix 64 × 64, 2 excitations. In total 240 volumes were collected through an 8-minute run.

#### Visit 2: Concurrent spTMS-fMRI

Prior to the TMS-fMRI runs, neuronavigation was conducted to localize stimulation targets for each participant with the Visor2 software (ANT Neuro, Enschede, NL) outside the scanner room. The right parietal stimulation site was defined as a key parietal node of the frontoparietal control network identified in previous studies ^19^. The right M1 was in the hand-knob region (**Table S4**). These targets were transformed from standard MNI space to individual’s native space and stereotaxically identified on their T1w images loaded in the Visor 2. The corresponding scalp stimulation locations were then marked on a Lycra swim cap (Speedo USA) worn by the participant into the scanner for TMS-fMRI runs. Immediately before scanning, the individual MT was defined in the MRI room as the stimulation intensity that induced visually observable hand motor activity in at least 5 of 10 consecutive trials. If no obvious hand movement was observed within 10 minutes of testing, stimulation was performed at 100% machine output.

During the TMS-fMRI runs, TMS pulses were delivered through an MRI-compatible air-cooled figure-eight TMS coil (MagPro MRI-B91) that was fixated by a custom-built holder and connected to a Magpro X100 stimulator (Magventure) in an adjoining room. Single pulses at 120% MT were presented with a jittered inter-stimulus interval ranging from 1-6 TRs to avoid the prediction of when TMS would be delivered. This rapid event-related design resulted in 68 stimulations per site over 167 volumes. For both stimulation sites, the TMS coil was perpendicular to the orientation of precentral gyrus (ca. 45° to the midline pointing in posterior direction) ^47^. The order of stimulation sites was randomized across participants.

The concurrent TMS-fMRI runs were acquired using the same scanner but with a TMS-compatible single-channel head coil that provides sufficient space for accommodating the TMS coil (**Fig. 2**). The TMS-fMRI images were acquired using a T2w gradient-echo spiral in/out pulse sequence: TE = 30 ms, flip angle = 85°, FOV = 22 cm, pixel size = 3.4 mm, 31 axial slices (4 mm thick) covering the whole brain, slice spacing 1 mm, matrix 64 × 64, 2 excitations. A 300 ms break was set between volumes (TA = 2100 ms) to allow single-pulse TMS delivery without affecting functional volume acquisition as we have done in our previous work ^19, 48^, resulting in an effective TR of 2400 ms. An automated high order shimming for spiral acquisitions was run prior to each TMS-fMRI run to reduce blurring and signal loss from field inhomogeneities.

Participants wore earplugs to protect their hearing and minimize the impact of sounds from TMS clicks and scanner noise. We recorded Subjective Units of Discomfort Scale (SUDS) immediately after each TMS-fMRI run. Participants were asked to report their pain/discomfort level from 0 to 100, with ‘0’ meaning ‘no pain’ and ‘100’ meaning ‘the worst pain you can imagine’. Heart rate (pulse oximeter clipped to the distal phalanx) and respiration (mid-thoracic strain gauge; both GE Healthcare) were acquired at 50 Hz during each fMRI run.

#### MRI processing

The heart rate and respiration during each fMRI run were processed offline and were regressed during data reconstruction where raw P-files were converted to NIFTI files. Preprocessing was performed for each fMRI run using the fMRIPrep pipeline ^36^ as that done in the concurrent TMS-iEEG experiment. The post-processing steps for rs-fMRI data were also the same as described for the TMS-iEEG experiment.

Following post-processing, resting-state functional connectivity (FC) was computed based on Fisher’s *z*-transformed Pearson correlation coefficient between BOLD times series extracted from the hippocampus and stimulation sites (parietal cortex and M1 respectively) for each participant. The stimulation masks were generated by a 6-mm radius GM sphere centered at the cortical stimulation locations in MNI space. Besides, we estimate the spatial proximity between the actual stimulation site and the individual optimal site defined by FC analyses. Specifically, we first identified the voxel exhibiting maximal FC to the hippocampus within a predefined anatomical mask either for the parietal cortex or M1. The parietal mask was generated by combining three parietal regions of A39rd, A40c, and A39rv from the Brainnetome (BN) atlas ^49^; The M1 mask was generated by combining two regions of A4hf and A4ul in the BN atlas. Euclidean distance was then calculated between this peak FC voxel and the actual stimulation site.

For TMS-fMRI data, post-processing included 0.008 Hz high-pass filtering and 6-mm FWHM Gaussian kernel smoothing. Next, first-level general linear model (GLM) analysis was performed for each participant with SPM 12 (https://www.fil.ion.ucl.ac.uk/spm/). Individual TMS delivery timepoints were convolved with canonical hemodynamic response function (HRF) with six head movement parameters as nuisance regressors. Following model estimation and contrast with implicit baseline, the average parameter estimates (*z*-scored beta weights) were extracted from the left or right hippocampus of each TMS-fMRI run for further statistical analyses.

#### Statistical analyses

One-sample *t*-tests (two-sided) were conducted to determine if TMS-evoked responses in the hippocampus were significantly greater or less than 0 when stimulating the parietal cortex or M1. To test whether individual variability in hippocampal responses could be explained by functional connectivity, we computed Pearson correlations between the hippocampal BOLD responses and the individual FC between the stimulation site and the hippocampus. In parallel, we examined whether spatial proximity contributed to response variability by correlating hippocampal responses with the Euclidean distance between the actual stimulation site and the individual’s FC-defined optimal parietal (or M1) site. For each computation, outliers defined as values exceeding 3 times SD from the mean of the group distribution were excluded to ensure stable and robust estimation.

### 3. Concurrent rTMS-iEEG Experiment

#### Participants

Four adult neurosurgical patients undergoing clinical intracranial electrode implantation for seizure monitoring were enrolled and met our inclusion criteria of having: 1) recording electrodes in the hippocampus, and 2) received parietal cortex rTMS. These patients were randomly assigned to the experimental group (2 patients) and the control group (2 patients). Three of the four participants were recruited from the concurrent spTMS-iEEG Experiment. Demographic information for all patients is provided in **Table S2**. All experimental procedures were approved by the University of Iowa Institutional Review Board (IRB). Written informed consent was obtained from all participants.

#### Experimental design

The experimental setup, including MRI imaging and iEEG recording procedures, was consistent with those used in the concurrent spTMS-iEEG experiment. The key difference was the TMS protocol. Each patient received 200 pulses of stimulation delivered at either 10 Hz or 20 Hz. These pulses were organized into five trains, with each lasting 30 secs. Each train began with a TMS-on period with the delivery of 40 pulses (10 Hz: 4 secs; 20 Hz: 2 secs) and then followed by a TMS-off period (10 Hz: 26 secs; 20 Hz: 28 secs). The MNI coordinates and stimulation frequency of rTMS for each patient are provided in **Table S3**. Sham stimulation was delivered by flipping the TMS coil 180°, and sham sessions always preceded active sessions to prevent potential carry-over effects. For participants who also completed the spTMS-iEEG protocol, the rTMS experiment was conducted after the spTMS experiment.

#### iEEG preprocessing

Preprocessing followed procedures similar to those used in the spTMS-iEEG experiment. First, the rTMS artifact window (−10 ms before train onset to 25 ms after train offset) was removed. Filters were then applied to the remaining signal, and the artifact window was interpolated. The resulting data were subsequently downsampled to 1000 Hz and segmented into epochs from −5 to 30 secs relative to train onset. No decay correction was applied here, in order to preserve potential lasting effects induced by rTMS. Besides, to minimize the influence of potential spikes occurring outside the stimulation period on subsequent time-frequency analysis, time points with amplitudes exceeding 5 times the SD of the entire epoch were identified, and ±10 ms around those points were interpolated.

#### Statistical analyses

The preprocessed data were then subjected to time-frequency analysis to assess the modulation effects on the hippocampal activities in various frequency bands by parietal rTMS. The time-frequency decomposition, by the same method as in Experiment 1, was first performed for each train to extract spectral power, which was then baseline-corrected by subtracting the average power of a time window [−1, −0.05 secs]. Spectral power was subsequently averaged within each frequency band, across the five stimulation trains, and within the 0–25 secs post-rTMS window. rTMS artifacts were unlikely to contaminate this window, given a sufficient interval (100 ms for 10 Hz; 50 ms for 20 Hz) from the onset of the final pulse in the rTMS train. One-sample *t*-tests (two-sided) were then conducted to evaluate power changes relative to the baseline. Finally, power values were compared between TMS conditions (active versus sham) and across TMS targeting strategies (Hippocampal-FC-guided versus non-hippocampal-FC-guided) to identify frequency bands significantly modulated by hippocampal-FC-guided active rTMS.

