## Supplementary information for "Multimodal Evidence for Hippocampal Engagement and Modulation by Functional Connectivity-Guided Parietal TMS"

**Table S1. iTEP amplitudes (SD) across different groups and TMS conditions.**
Significant results are indicated by asterisks (*, $p_{FDR}$ < .05).

| | Time window | Active TMS | | | Sham TMS | | | Active – Sham (contrast) | | |
|---|---|---|---|---|---|---|---|---|---|---|
| | | Mean ± s.d. | *t* | $p_{FDR}$ | Mean ± s.d. | *t* | $p_{FDR}$ | Mean ± s.d. | *t* | $p_{FDR}$ |
| **Hippocampal-FC-guided TMS; Responsive contacts** | 26-49 ms | -2.46 ± 0.71 | -10.42 | **4.22E-05*** | -0.20 ± 0.71 | -0.87 | 0.478 | -2.26 ± 0.31 | -22.06 | **5.09E-07*** |
| | 108-175 ms | 3.87 ± 1.22 | 9.55 | **6.45E-05*** | 0.37 ± 0.47 | 2.37 | 0.087 | 3.50 ± 0.99 | 10.61 | **4.22E-05*** |
| | 224-334 ms | -3.00 ± 1.27 | -7.08 | **3.13E-04*** | -0.28 ± 0.29 | -2.90 | **0.045*** | -2.72 ± 1.08 | -7.58 | **2.80E-04*** |
| **Hippocampal-FC-guided TMS; Non-responsive contacts** | 26-49 ms | -0.72 ± 1.09 | -2.19 | 0.097 | -1.27 ± 0.65 | -6.46 | **2.80E-04*** | 0.55 ± 1.58 | 1.15 | 0.354 |
| | 108-175 ms | 1.60 ± 1.17 | 4.54 | **0.003*** | 1.16 ± 1.91 | 2.01 | 0.115 | 0.44 ± 1.83 | 0.80 | 0.478 |
| | 224-334 ms | -0.12 ± 1.24 | -0.31 | 0.793 | 0.23 ± 0.50 | 1.54 | 0.233 | -0.35 ± 1.10 | -1.05 | 0.388 |
| **Non-Hippocampal-FC-guided TMS** | 26-49 ms | -0.92 ± 0.70 | -5.23 | **3.13E-04*** | -2.52 ± 1.40 | -7.21 | **4.07E-05*** | 1.61 ± 1.61 | 3.99 | **0.003*** |
| | 108-175 ms | 0.37 ± 0.64 | 2.32 | 0.072 | 0.91 ± 2.62 | 1.39 | 0.248 | -0.54 ± 2.76 | -0.79 | 0.478 |
| | 224-334 ms | 0.00 ± 1.08 | 0.01 | 0.994 | -0.64 ± 1.83 | -1.41 | 0.248 | 0.65 ± 1.30 | 1.99 | 0.110 |

#### Table S2. Patient demographic information in Exp 1&3

| Patient ID | Recruitment | Age | Sex | Handedness | Ethnicity | Education (yr) | Age of onset (yr) | Onset of Epilepsy | Lesions | Prior surgical resection |
|---|---|---|---|---|---|---|---|---|---|---|
| 403 | Exp 1 | 56 | F | R | non-Hispanic White | 15 | 19 | left anterior temporal | none | No |
| 405 | Exp 1&3 | 19 | M | R | non-Hispanic White | 12 | 9 | left frontal | left frontal | No |
| 416 | Exp 1&3 | 34 | M | - | non-Hispanic White | 12 | 20 | left occipital | focal cortical dysplasia type IIb | No |
| 423 | Exp 1&3 | 51 | M | R | non-Hispanic White | 8 | * | left anterior temporal | none | No |
| 430 | Exp 1 | 28 | M | R | non-Hispanic White | 12 | 4 | generalized | none | No |
| 477 | Exp 1 | 23 | F | R | non-Hispanic White | 12 | 16 | left lateral posterior parietal | none | No |
| 493 | Exp 3 | 35 | M | L | non-Hispanic White | 16 | 7 | right medial temporal & orbitofrontal | none | No |
| 820 | Exp 1 | 51 | M | R | non-Hispanic White | 16 | 36 | left temporal | none | No |

### Table S3. TMS-iEEG experiment information in Exp 1&3

| Patient ID | TMS strategy | MNI coordinates of TMS Site | Exp1 (spTMS-iEEG) | Exp3 (rTMS-iEEG) | MT (% machine output) | Stimulation Intensity (% MT) | rTMS stimulation frequency | # Electrodes (# hippocampus) | Implantation Type | Day of testing medications |
|---|---|---|---|---|---|---|---|---|---|---|
| 403 | Non-Hippo-FC-guided | 12, -86, 42 | Yes | - | 70% | 100% | - | 231 (11) | ECoG & sEEG | biotin, cholecalciferol, clonazepam, diazepam, fluoxetine HCl, hydroxyzine HCl, lamotrigine, levothyroxine sodium, magnesium oxide, metoclopramide HCL, oxcarbazepine, perampanel |
| 405 | Non-Hippo-FC-guided | 39, -80, 41 | Yes | Yes | 88% | 100% | 20 Hz | 170 (1) | ECoG & sEEG | acetaminophen, bisacodyl, cefazolin, docusate, heparin, levetiracetam, Lorazepam, magnesium hydroxide, morphine, omeprazole, ondansetron, oxcarbazepine, perampanel, sennosides, sodium chloride 0.9% |
| 416 | Hippo-FC-guided | 24, -61, 72 | Yes | Yes | 59% | 100% | 10 Hz | 192 (9) | ECoG & sEEG | lacosamide, levetiracetam, cannabidiol oil |
| 423 | Hippo-FC-guided | 32, -67, 64 | Yes | Yes | 49% | 100% | 20 Hz | 213 (9) | ECoG & sEEG | albuterol sulfate, carbamazepine, fluticasone, propionate, guaifenesin/dextromethorphan, ibuprofen, mirtazapine, tiotropium bromide, zonisamide |
| 430 | Non-Hippo-FC-guided | -6, -33, 78 | Yes | - | 53% | 100% | - | 72 (1) | sEEG | clonazepam, lamotrigine, risperidone, topiramate, venlafaxine HCl |
| 477 | Hippo-FC-guided | -32, -85, 43 | Yes | - | 59% | 120% | - | 148 (3) | ECoG & sEEG | albuterol sulfate, clonazepam, fluticasone propion/salmeterol, ipratropium/albuterol sulfate, levetiracetam, montelukast sodium, oxcarbazepine, sertraline HCl, topiramate |
| 493 | Non-Hippo-FC-guided | 55, -50, 53 | - | Yes | 69% | 120% | 10 Hz | 222(5) | ECoG & sEEG | levetiracetam, lamotrigine and oxcarbazepine |
| 820 | Non-Hippo-FC-guided | -59, -49, 46 | Yes | - | 62% | 100% | - | 230(3) | ECoG & sEEG | omeprazole, propranolol |

**Table S4. Participants' demographic information in Exp 2 (TMS-fMRI)**

Seventy-nine neurologically healthy participants were recruited. Of these, 50 participants received TMS to the right inferior parietal lobe (IPL); 78 participants received TMS to the right primary motor cortex (M1), which served as a control site. All participants underwent resting-state fMRI scanning.

| TMS-fMRI/rs-fMRI | No. of participants | Female | Male | Age (s.d.) | Years of Education (s.d.) | MNI coordinates for TMS |
|---|---|---|---|---|---|---|
| R M1 | 78 | 48 | 30 | 31.4 (10.3) | 16.1 (2.2) | 40, -18, 64 |
| R IPL | 50 | 30 | 20 | 31.8 (11.3) | 16.1 (2.2) | 48, -54, -46 |
| Rs-fMRI | 79 | 48 | 31 | 31.6 (10.4) | 16.1 (2.1) | / |

**Figure S1. The parietal TMS sites in Exp 1 & 3.**
Screenshots from the Brainsight Neuronavigation software (Rogue Research, Montreal, Quebec, Canada) display the TMS target locations (marked by the blue dots) for each subject. For the Hippocampal-FC-guided group, the individual rsFC map was loaded to identify the parietal spot exhibiting the maximal functional connectivity to the hippocampus.

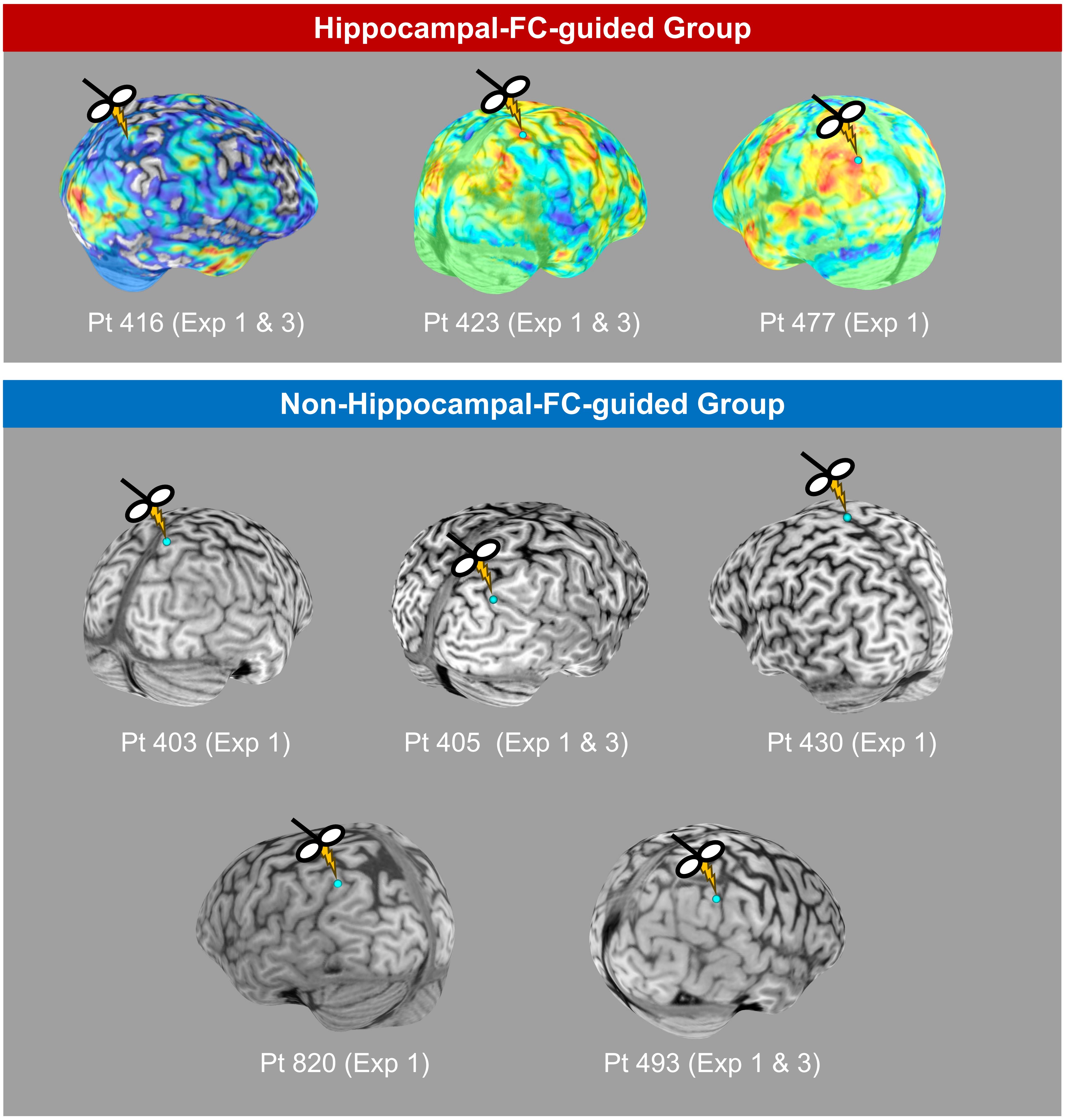

**Figure S2. Location of TMS sites, hippocampus contact, and individual iTEPs.**
iTEP waveforms of each hippocampal contact are shown. Colored lines represent the iTEPs by active TMS; gray lines represent the iTEPs by sham TMS. Different colors indicate different patients. Responsive contacts to active parietal TMS are indicated by symbols: crosses (+) mark contacts with iTEP amplitudes exceeding 5 standard deviations relative to baseline in the active condition but not in the sham condition; asterisks (*) mark contacts showing significant differences between active and sham conditions based on point-by-point comparisons ($p_{FDR} < .05$).

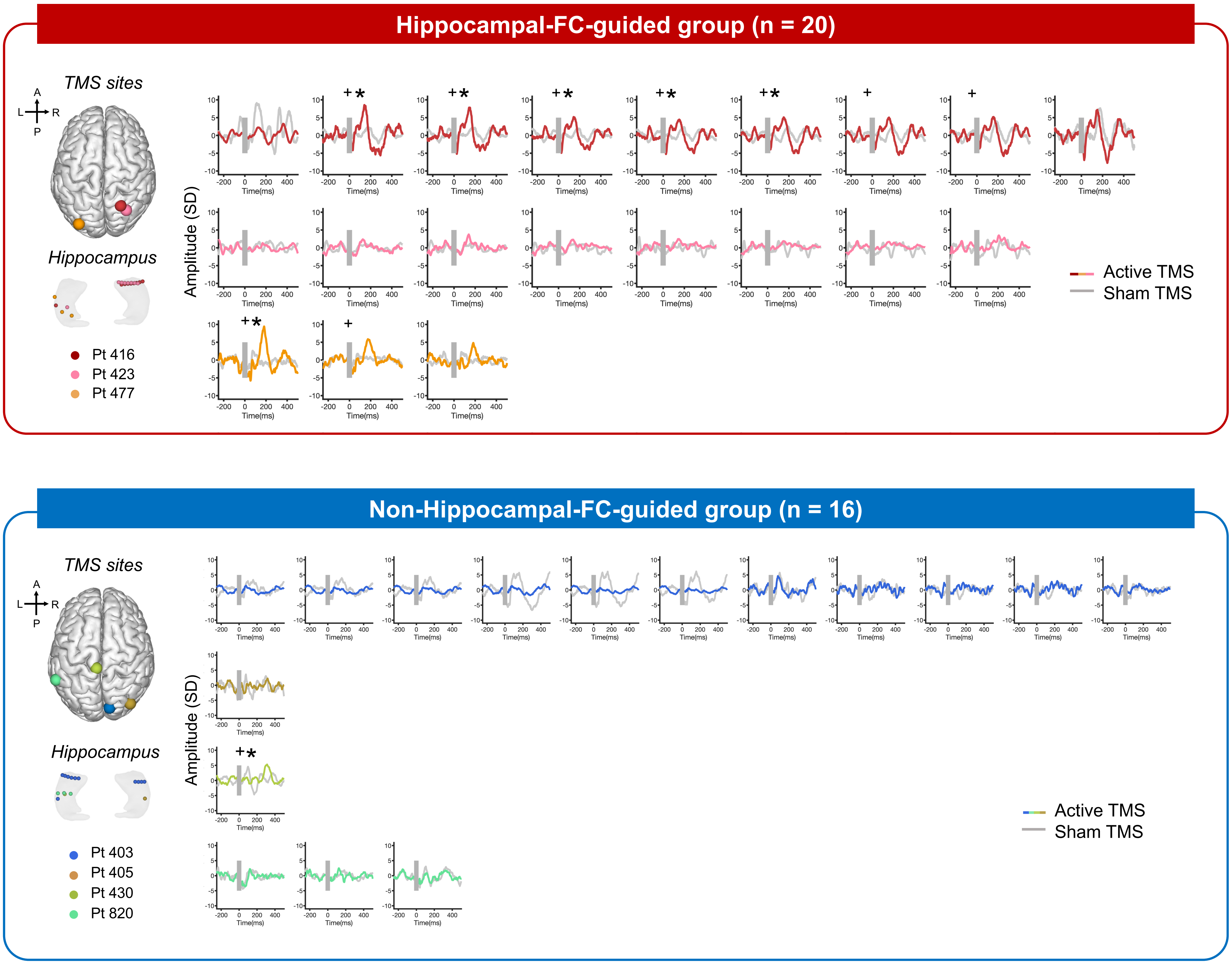

**Figure S3. Hippocampal modulation following each train of Hippocampal-FC-guided active rTMS.**
Asterisks (*) indicate significant time bins ($p_{FDR} < .05$).

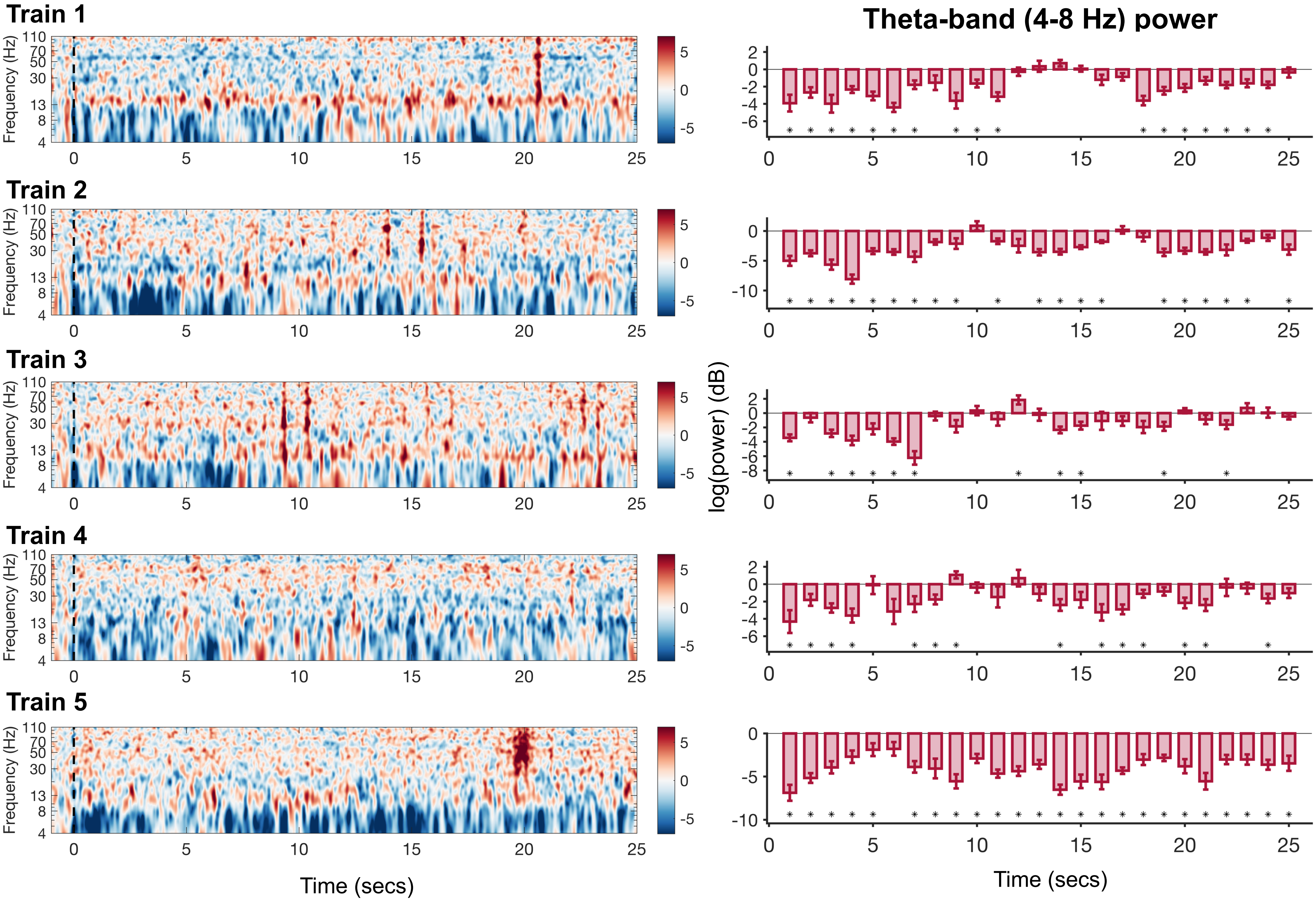

**Figure S4. Parahippocampal and Amygdala modulation by Hippocampal-FC-guided parietal rTMS**

(A) Spectral power changes in the parahippocampal following Hippocampal-FC-guided active rTMS.

(B) Spectral power changes in the amygdala following Hippocampal-FC-guided active rTMS.

(C) Comparison of theta power changes across different regions. Significant theta power suppression was only observed in the hippocampus ($p_{FDR} < .001$), but not in the parahippocampal gyrus or amygdala ($p_{FDR} = .07, .33$). Besides, the theta power suppression in the hippocampus was significantly greater than that in the parahippocampal gyrus and amygdala ($p_{FDR} = .045, .004$).
Hippo = hippocampus; PrHippo = parahippocampal gyrus; Amyg = amygdala.

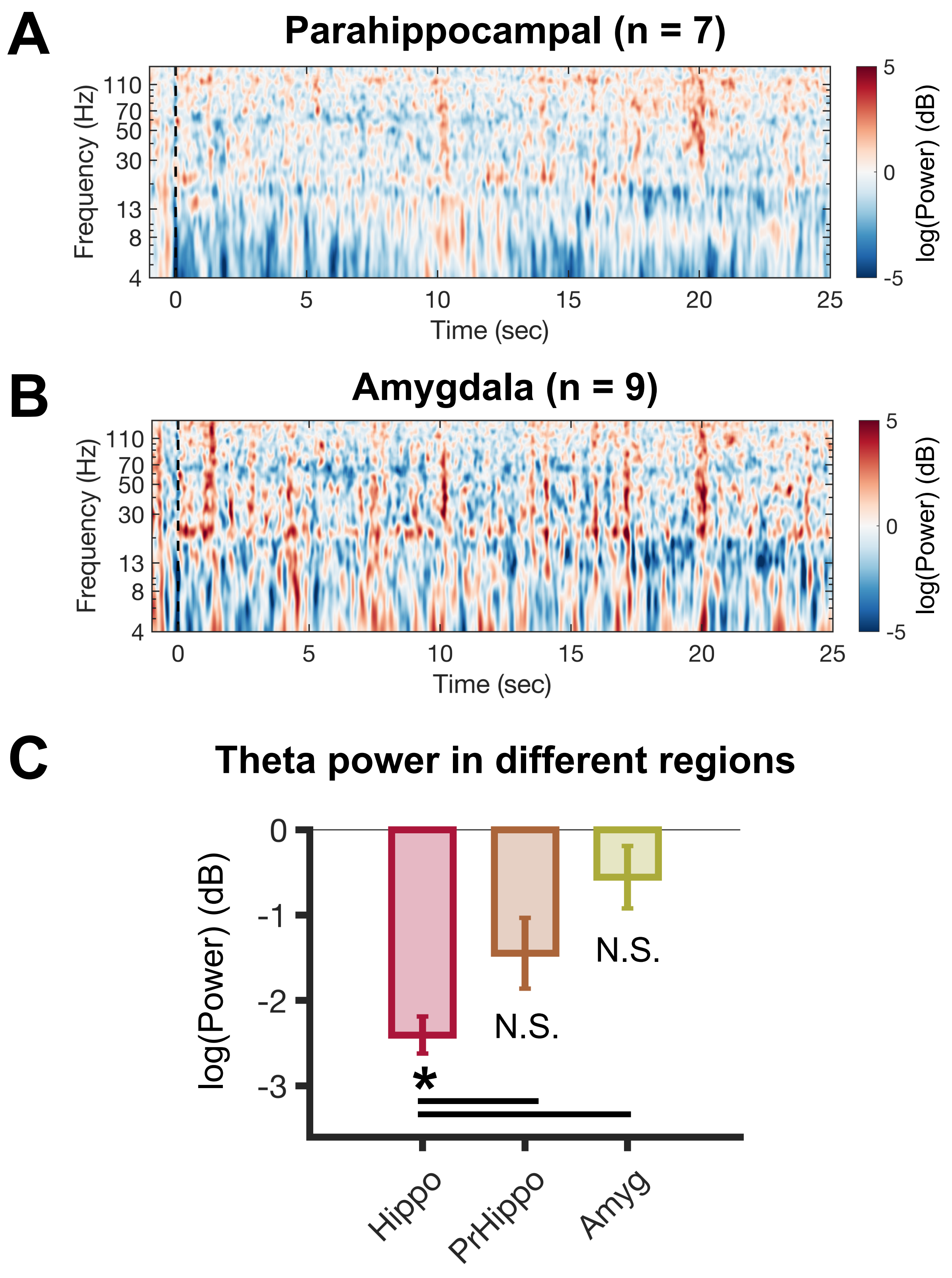
